# Improving T-cell mediated immunogenic epitope identification via machine learning: the neoIM model

**DOI:** 10.1101/2022.06.03.494687

**Authors:** Lena Pfitzer, Lien Lybaert, Cedric Bogaert, Bruno Fant

## Abstract

The identification of immunogenic peptides that will elicit a CD8+ T cell-specific immune response is a critical step for various immunotherapeutic strategies such as cancer vaccines. Significant research effort has been directed towards predicting whether a peptide is presented on class I major histocompatibility complex (MHC I) molecules. However, only a small fraction of the peptides predicted to bind to MHC I turn out to be immunogenic. Prediction of immunogenicity, i.e. the likelihood for CD8+ T cells to recognize and react to a peptide presented on MHC I, is of high interest to reduce validation costs, de-risk clinical studies and increase therapeutic efficacy especially in a personalized setting where *in vitro* immunogenicity pre-screening is not possible.

To address this, we present neoIM, a random forest classifier specifically trained to classify short peptides as immunogenic or non-immunogenic. This first-in-class algorithm was trained using a positive dataset of more than 8000 non-self immunogenic peptide sequences, and a negative dataset consisting of MHC I-presented peptides with one or two mismatches to the human proteome for a closer resemblance to a background of mutated but non-immunogenic peptides. Peptide features were constructed by performing principal component analysis on amino acid physicochemical properties and stringing together the values of the ten main principal components for each amino acid in the peptide, combined with a set of peptide-wide properties. The neoIM algorithm outperforms the currently publicly available methods and is able to predict peptide immunogenicity with high accuracy (AUC=0.88). neoIM is MHC-allele agnostic, and *in vitro* validation through ELISPOT experiments on 33 cancer-derived neoantigens have confirmed its predictive power, showing that 71% of all immunogenic peptides are contained within the top 30% of neoIM predictions and all immunogenic peptides were included when selecting the top 55% of peptides with the highest neoIM score. Finally, neoIM results can help to better predict the response to checkpoint inhibition therapy, especially in low TMB tumors, by focusing on the number of immunogenic variants in a tumor.

Overall, neoIM enables significantly improved identification of immunogenic peptides allowing the development of more potent vaccines and providing new insights into the characteristics of immunogenic peptides.

## Introduction

The goal of cancer immunotherapy is to activate a patient’s own immune system to very specifically recognize and kill tumor cells. In this framework, a tumor cell is recognized by CD8+ cytolytic T lymphocytes by inspecting the cell’s presented antigens derived from mutated genome sequences^1^. These tumor-specific mutated peptides, also called neoantigens, play a crucial role in targeted cancer therapies like personalized neoantigen vaccines or adoptive T cell therapy as well as in treatment with immune checkpoint inhibitors.

While encouraging, clinical results from earlier trials with immunotherapeutic cancer vaccines have remained modest, leading to doubt about the efficacy of this form of immunotherapy. However, many trials’ low efficacy can be attributed to the targeting of tumor associated antigens (TAAs) rather than neoantigens. These TAAs are self-proteins that are over-expressed in cancer cells and shared by many patients. Developing T cells that do recognize such self-antigens are subjected to tolerance induction mechanisms in the thymus to avoid auto-immunity. This might be a reason why it appears to be rather difficult to induce high-avidity T cells recognizing tumor cells expressing TAAs^2,3^. Other reasons for limited clinical efficacy of cancer vaccines may include the suboptimal activity of the currently used vaccine platforms, tumor heterogeneity and the immunosuppressive tumor microenvironment (TME) as well as other issues such as target cancer patients’ stratification and clinical trial design^4^.

More recent studies have shown that it is the neoantigens, which are exclusively expressed by tumor cells, that drive the response to CTLA-4 and PD-1 blockade^5–7^. Additionally, acquired resistance to therapy has been linked to loss of neoantigen expression, in non-small-cell lung cancer patients for example^8^. Personalized neoantigen vaccines have been shown to successfully activate neoantigen-specific T cells, resulting in clinical responses^9–11^. This highlights the important role of neoantigens and the relevance of personalized or neoantigen-targeting therapies. It is therefore crucial to correctly identify neoantigens that are able to evoke an immune response as only a minor fraction of neoantigens is truly immunogenic. In a recent consortium study however, only 6% of the top neoantigens predicted by multiple academic, clinical, and commercial teams could be validated as immunogenic^12^. This is consistent with other studies in which only a low percentage of predicted immunogenic neoantigens can actually be validated and highlights the challenge in identifying truly immunogenic material^13–15^.

Enzyme-linked immunosorbent spot (ELISPOT) assays are currently the most often used method to measure the presence of neoantigen-specific T cells ^16^. Whilst ELISPOT can be a useful tool in neoepitope validation, the necessary cost and time investment limits its feasibility, especially in the context personalized therapy manufacture, where time constraints are highly stringent. Accurate immunogenicity prediction through computational methods is therefore of high importance to de-risk clinical studies and increase therapeutic efficacy.

MHC-I binding affinity of a peptide is currently assumed to be one of the most important factors of neoantigen immunogenicity, hence many neoantigen prediction workflows use it as the key parameter for prioritization^17^. However, peptide presentation alone does not guarantee an immune response since, as mentioned earlier, only a minor fraction of peptides shown to be presented result in a positive response^13,18^. Additional metrics have been suggested to more accurately predict peptide immunogenicity, such as the differential agretopicity index (DAI) which is the ratio of MHC binding affinity between the mutated and corresponding normal peptide^19–21^, the expression level of the mutated gene, the peptide-MHC binding stability, the peptide hydrophobicity, the dissimilarity to the proteome or, conversely, the similarity to known bacterial or viral antigens, and the position of the mutation in the peptide^21–26^. This variety in evaluation metrics indicates that there are numerous criteria that influence the immunogenicity of presented peptides. So far however, it is still unclear what the exact properties are that lead to recognition by CD8+ T cells and a subsequent immune reaction.

Even though thousands of immunogenic peptides have been identified through various types of assays and collected in the immune epitope database (IEDB)^27^, methods for accurate *in silico* prediction of peptide immunogenicity are still lacking. A major problem is the lack of consistent data on neoantigen immunogenicity across different individuals, since due to the high level of MHC polymorphisms and individual immune contexture, a neoantigen not being able to activate one individuals’ T cells can still elicit a response in other individuals. It is clear that a strongly immunogenic peptide should be able to cause an immune response in a large proportion of the population presenting the peptide. However, a peptide that is inherently immunogenic might not be presented by the HLA alleles of an individual, making it invisible to their immune system. Additionally, even if a peptide is immunogenic in a large proportion of the population, due to the individualized nature immune repertoire, not every patient might generate an immune response against that peptide. This is represented in studies, where some patients develop an immune response towards specific peptides and some do not^28,29^. Therefore, it is difficult to assess if a certain peptide is truly non-immunogenic.

In our study, we introduce a novel method to assemble a non-immunogenic dataset, by using peptides that have a similar but not identical sequence to the human proteome. Subsequently, we present neoIM, a first-in-class algorithm trained to predict the immunogenicity of MHC-I presented peptides. We show that this method is able to significantly reduce the number of peptides falsely classified as immunogenic based on MHC presentation prediction alone. The IEDB immunogenicity predictor, the only other comparable tool, is strongly outperformed by neoIM. *In vitro* validation was performed through ELISPOT experiments on cancer-derived neoantigens which were all predicted to be presented, confirming that further prioritization with neoIM significantly increases the true positive rate. Furthermore, in low TMB melanomas, the number of immunogenic variants predicted by neoIM was able to better inform on the response to checkpoint inhibitor therapy than the general tumor mutational burden. Therefore, prioritizing neoepitopes with high predicted immunogenicity by neoIM increases the likelihood of selecting clinically actionable neoantigens.

## Methods

The flow chart for neoIM’s training and prediction workflow is shown in Figure 1.

**Figure 1:**
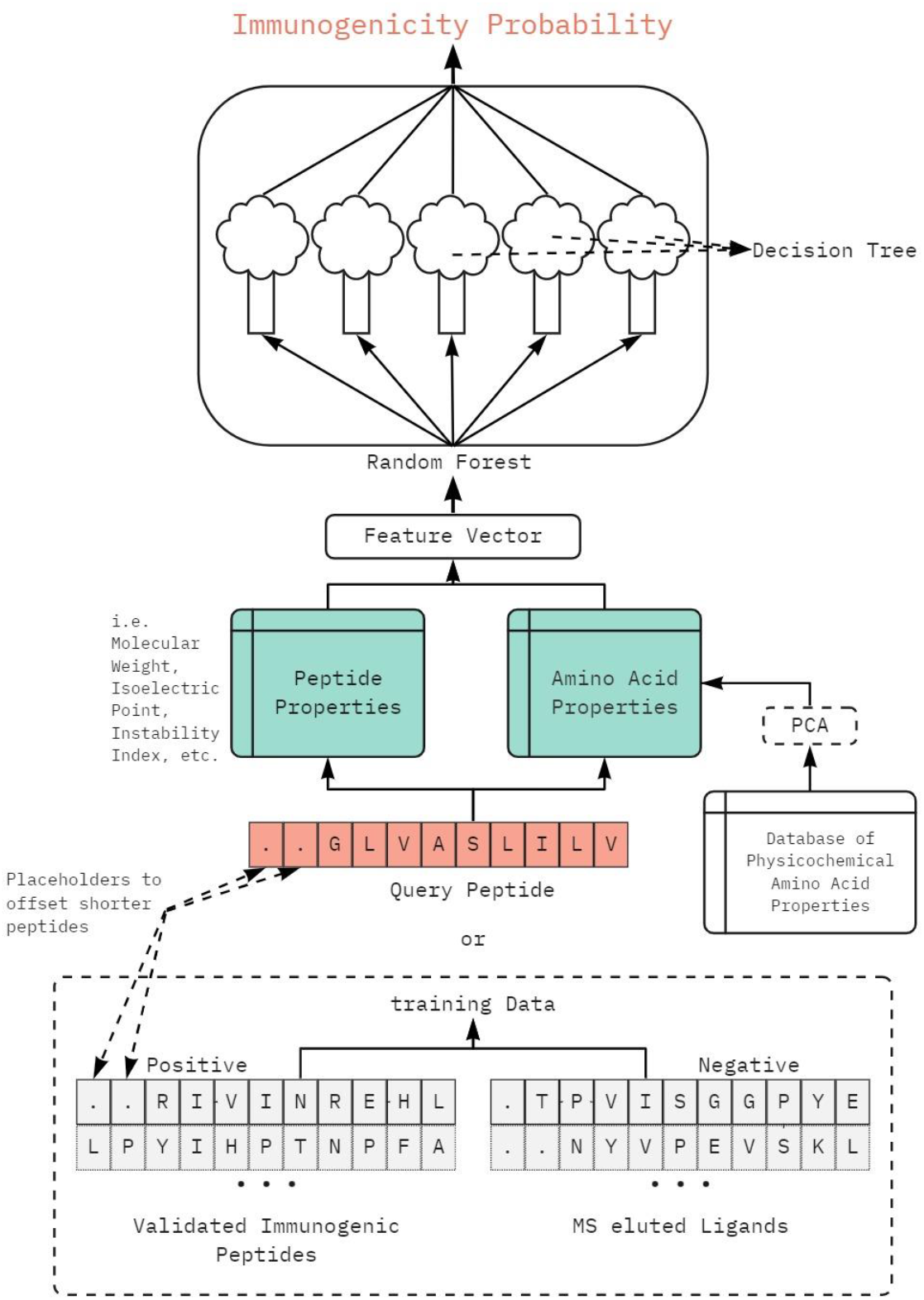
Flowchart of neoIM’s training and prediction workflow.

### Training Dataset

#### Positive training dataset

The positive dataset was mainly composed of immunogenic peptides collected from the IEDB database (version June. 01, 2021^27^). A peptide was considered positive if it was a linear sequence, tested for *Homo sapiens*, resulting in at least one positive immunogenic assay (Error! Reference source not found.) and being of length 9, 10 or 11 amino acids. Additional sources of positive peptides are listed in Error! Reference source not found..

In a second filtering step, every peptide was blasted against the human proteome. Self-peptides, defined as perfect matches to the human proteome (UNIPROT ID: UP000005640), were removed from the dataset. Duplicate peptides were filtered out. Positive data after such filtering steps comprised 8258 nona-, deca- and undecamers (9-11mers).

#### Negative training dataset

Non-immunogenic peptides were collected from IEDB data of MHC binding peptides (version Feb. 19, 2020^27^). Peptides were filtered to be a linear sequence, from *Homo sapiens*, being of length 9, 10 or 11 amino acids and validated as MHC binders to a human. The peptides were also blasted against the human proteome. Only peptides that were a close but imperfect match to the human proteome (1 to 3 mismatches) were retained. Duplicate peptides and peptides present in the positive training dataset were filtered out. Negative data after such filtering steps comprised 6872 nona-, deca- and undecamers. A χ2 Goodness-Of-Fit Test was performed to assess whether the peptide length distributions are significantly different between the positive and negative dataset.

### Peptide encoding

To train a model on our dataset, peptides were encoded in a numerical sequence. For this purpose, amino acid physicochemical properties were retrieved from the AAindex1 database^30^. The database contains 553 properties for the 20 proteinogenic amino acids. After dropping 13 properties with missing values, the resulting 20×540 matrix was scaled to Z-values using the sklearn StandardScaler module. To reduce the dimensionality and noise of the data, principal component analysis was performed on the scaled feature matrix. Each amino acid was then translated into a value vector comprising its first ten principal components (PCs), which explain 89% of the total variance (Supplementary Figure 1B). Each peptide was then encoded by an ordered concatenation of the value vectors derived from each amino acid. To align peptides of different length (9-11aa) at the C-terminus, missing positions were imputed at the beginning as zeros as the PC values are distributed around zero. Some additional metrics at the peptide level were added using the biopython Bio.SeqUtils.ProtParam ^31^ module: molecular weight, aromaticity, instability index, isoelectric point, gravy index and secondary structure metrics (fraction of amino acids that tend to be in helix, turn or sheet). This resulted in a feature vector of length 118 for each peptide.

### Algorithm parameters and training

To create neoIM, a Random Forest model was trained on the assembled dataset using the scikit-learn ensemble module RandomForestClassifier^32^, using 10-fold cross-validation to optimize the model’s parameters. For the final model, the number of trees was fixed at ten thousand, information gain was used to measure the quality of a split, the minimum number of samples required for a split was 3 and no bootstrap samples were used when building trees. Feature importance was extracted from the RandomForestClassifier.

### Datasets for bias testing

One hundred thousand peptides were randomly sampled in equal proportions of length 9, 10 and 11 amino acids from the human proteome. MHC binding affinity was predicted using MHCnuggets^33^ for 38 HLA-A alleles, 52 HLA-B alleles and 12 HLA-C alleles. For each HLA allele, the 1% of peptides with the strongest binding affinity were considered presented peptides, which equates to a Rank (%) score^34^ < 1, and the neoIM immunogenicity score calculated. To investigate a potential bias towards certain HLA alleles, the fractions of peptides called as immunogenic (score >= 0.64) for each HLA allele were compared. To investigate a potential bias towards certain peptide lengths, the fractions of peptides called as immunogenic (score >= 0.64) for each length (9, 10 and 11 amino acids) were compared.

### Benchmarking

The neoIM algorithm was compared to the IEDB immunogenicity predictor, a model that rates peptides based on the position and properties of their amino acids^35^. The data included in the benchmarking dataset is described in Table 1.

**Table 1:** Sources of peptides included in the benchmarking dataset

| Year of publication | PMID | Source | Non-immunogenic epitopes | Immunogenic epitopes |
| --- | --- | --- | --- | --- |
| 2011 | 21918184 | Weiskopf et al.<sup>36</sup> | 477 | 42 |
| 2018 | 29397015 | Luxenburger et al.<sup>37</sup> | 100 | 26 |

Peptides classified as non-immunogenic in these datasets were screened against the IEDB database to remove potential false negatives, i.e., peptides that were recorded to result in a positive immunogenicity assay in another setting. Furthermore, only peptides predicted to be presented according to previously published affinity predictions^38^ were used for the benchmarking (NetMHCpan 4.0. Rank (%) < 2^34^). This resulted in overall 67 positives peptides and 473 negative peptides. Benchmark peptides which were also present in the training dataset, were removed from the training dataset and the algorithm retrained for benchmarking.

### Survival analysis of cohort treated with CPI

In a cohort of 64 melanoma patients treated with CPI therapy, different biomarkers were extracted for comparison purposes ^39^. First, the number of mutations per patient was extracted from the genomic variation data. Next, the number of peptide-presenting non-synonymous mutations was computed using MHCnuggets^33^. A mutation was considered peptide-presenting if it generated at least one peptide with rank (%) < 2^34^. Finally, the number of strongly immunogenic mutations was derived from predictions output by the neoIM algorithm. A mutation was considered strongly immunogenic if it generated at least one presented epitope (MHCnuggets^33^ rank (%) <2) with a neoIM score > 0.7. Survival analysis was performed to estimate probabilities of patients responding to therapy relative to the biomarkers. The cohort was first separated into high TMB and low TMB patients, using the 100 mutations per exome threshold commonly reported for melanoma studies ^39–41^. For either the full cohort, TMB-low patients or TMB-high patients, members were further divided at the median value of the biomarker to get two evenly sized groups, and their Kaplan-Meier curves were compared. Statistical testing was performed using the logrank test.

### In vitro validation

Peptides for *in vitro* validation were selected from Wells et al.^12^, from mutations identified in privately sequenced melanoma patients or from recurrently observed mutation events in certain cancer indications based upon data generated by the TCGA Research Network: https://www.cancer.gov/tcga. The selected peptides were predicted to be presented (MHCnuggets^33^ rank (%) <2) on at least two healthy donor HLA alleles (Error! Reference source not found.).

Experimental immunogenicity testing of the 33 selected peptides was performed by ELISPOT assay on healthy donor T-cells. Assays were performed with two rounds of T cell stimulation with autologous dendritic cells (DCs) loaded with peptides. All tests were performed in duplicate or triplicate. All peptides were predicted to be presented on at least two donor HLA alleles using netMHCpan4.0. A positive ELISPOT result was defined as a minimum of 25 spots per 5×10^4^ cells and an increase of at least 2x compared to the control (autologous DCs loaded with DMSO).

For the ROC and lift curves, predictions for each peptide by the neoIM algorithm were compared to the IEDB immunogenicity predictor and NetMHCpan 4.0^34^. NetMHCpan was included in the comparison in two different ways. First the rank score for HLA-A*02:01, which was shared by all donors, and second, the lowest rank score of all donor HLA alleles.

To assess if the scores of positive and negative peptides in the *in vitro* validation dataset are significantly different for various scoring methods, independent t-tests were performed.

## Results

### Building the training dataset

Training data was collected from multiple databases with the aim to resemble a mutanome, which consists mostly of mutations that are close matches to the proteome, only differing in a few positions. Therefore, all peptides in the training dataset should be resulting from genetic variation, be presented on MHC and only differ in their immunogenicity. A total of 8258 validated immunogenic epitopes with at least one mismatch to the human proteome comprised the positive training dataset. A negative training dataset was built from validated MHC binders that are a close but not perfect match to the human proteome (1 to 3 mismatches). This resulted in a negative training dataset of 6872 epitopes. The epitopes in the training dataset are between 9 and 11 amino acids in length. The detailed length distributions are shown in Supplementary Figure 1A.

The peptide length distribution between the positive and negative training dataset is slightly shifted with more 10- and 11mers in the positive dataset (26%, 9.2%) than in the negative dataset (17%, 6.4%) and less 9mers in the positive dataset (64.7%) than in the negative dataset (76.6%) (p-value = 0.02).

To get better insights into the characteristics of immunogenic peptides, the differential amino acid usage between epitopes in the positive and negative datasets was investigated (figure2A): immunogenic peptides tend to contain more hydrophobic amino acids, especially at peptide positions 2 and 9 where Leucin and Valin are preferentially used. Amino acids in immunogenic peptides also more often contain sulphur or aromatic groups with aliphatic (non-polar and hydrophobic) side chains dominating at positions 2 and 9 (Figure 2B). Generally, immunogenic peptides are also enriched in neutrally charged and larger amino acids (Figure 2C, D). The preference for Leucin and Valine at positions 2 and 9, which are important anchor positions for MHC binding, could also indicate a bias for HLA-A*02:01 presented peptides in the positive training dataset^42^ (see also below in section “Training the model”). HLA-A*02:01 is the most prevalent HLA allele in the human population and the most used allotype for immunogenicity experiments (Error! Reference source not found.).

**Figure 2:**
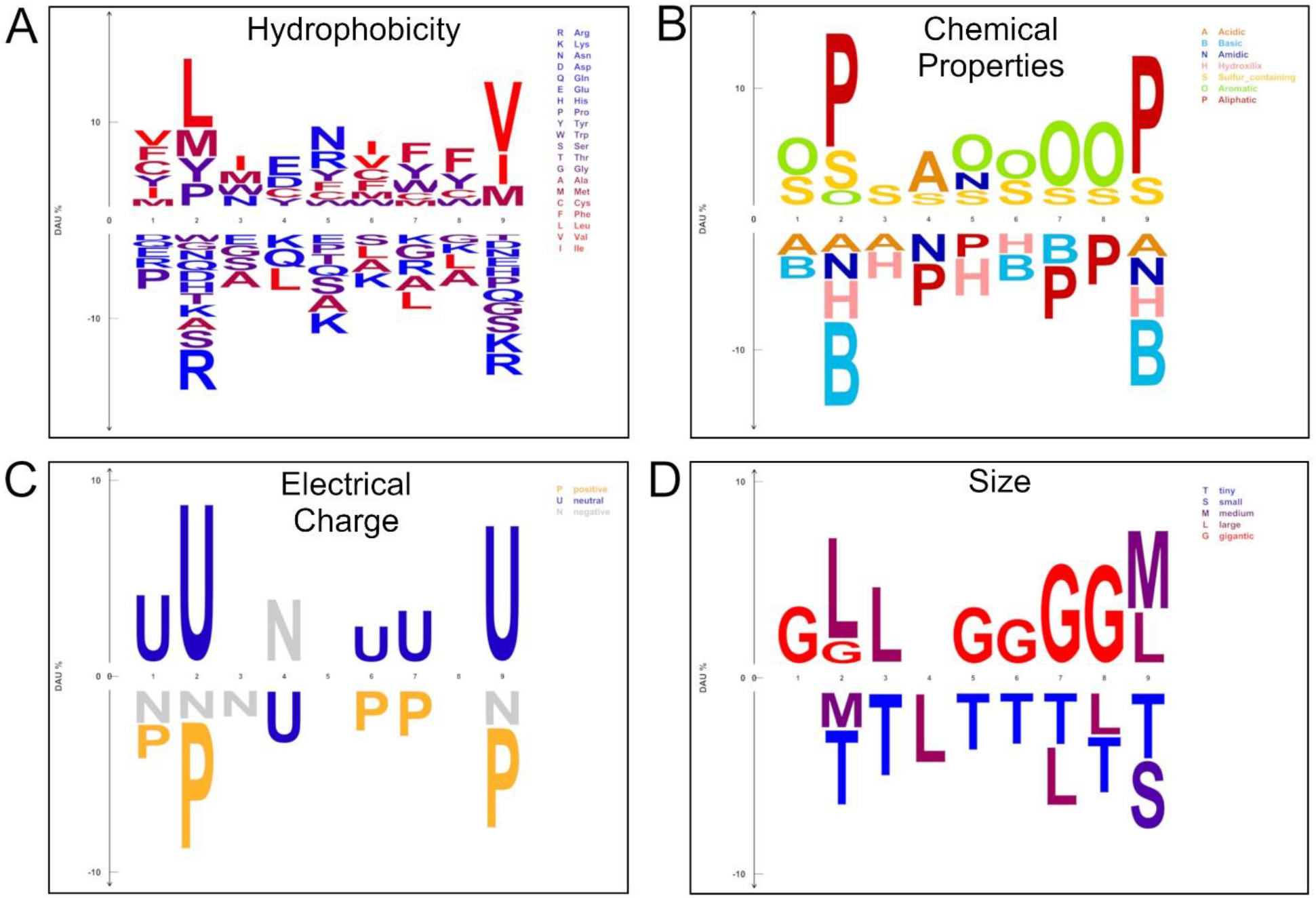
Logos showing differential amino acid usage of 9aa long peptides in the immunogenic positive dataset (top) over 9aa long peptides in the non-immunogenic negative dataset (bottom). Only peptides of length 9aa were analyzed. Fisher’s exact test was performed with a significance level of 0.05. Amino acids were either colored by hydrophobicity (A), grouped by chemical properties (B), electrical charge (C) or size (D).

### Training the model

By using a set of peptides all known to be presented by at least one MHC type, an HLA-agnostic model can be created that learns to identify immunogenic peptides based on their physicochemical properties alone. Amino acid properties from the AAindex1 database were retrieved and used to extract the 10 first principal components describing each amino acid across their physico-chemical properties. Each peptide was then translated into a numerical vector representing the amino acid sequence. Additionally, peptide-wide properties were added to the feature vector, like weight, isoelectric point, and hydrophobicity index. A random forest classifier was trained on the vectorized data, and its performance was assessed by ten-fold cross validation. The final model, dubbed neoIM, showed a consistently high performance with an average area under the Receiver Operator Characteristic (ROC) curve of 0.85 (Figure 3A) and an average precision (AP) of 0.91 for the precision recall curve (Figure 3B).

**Figure 3:**
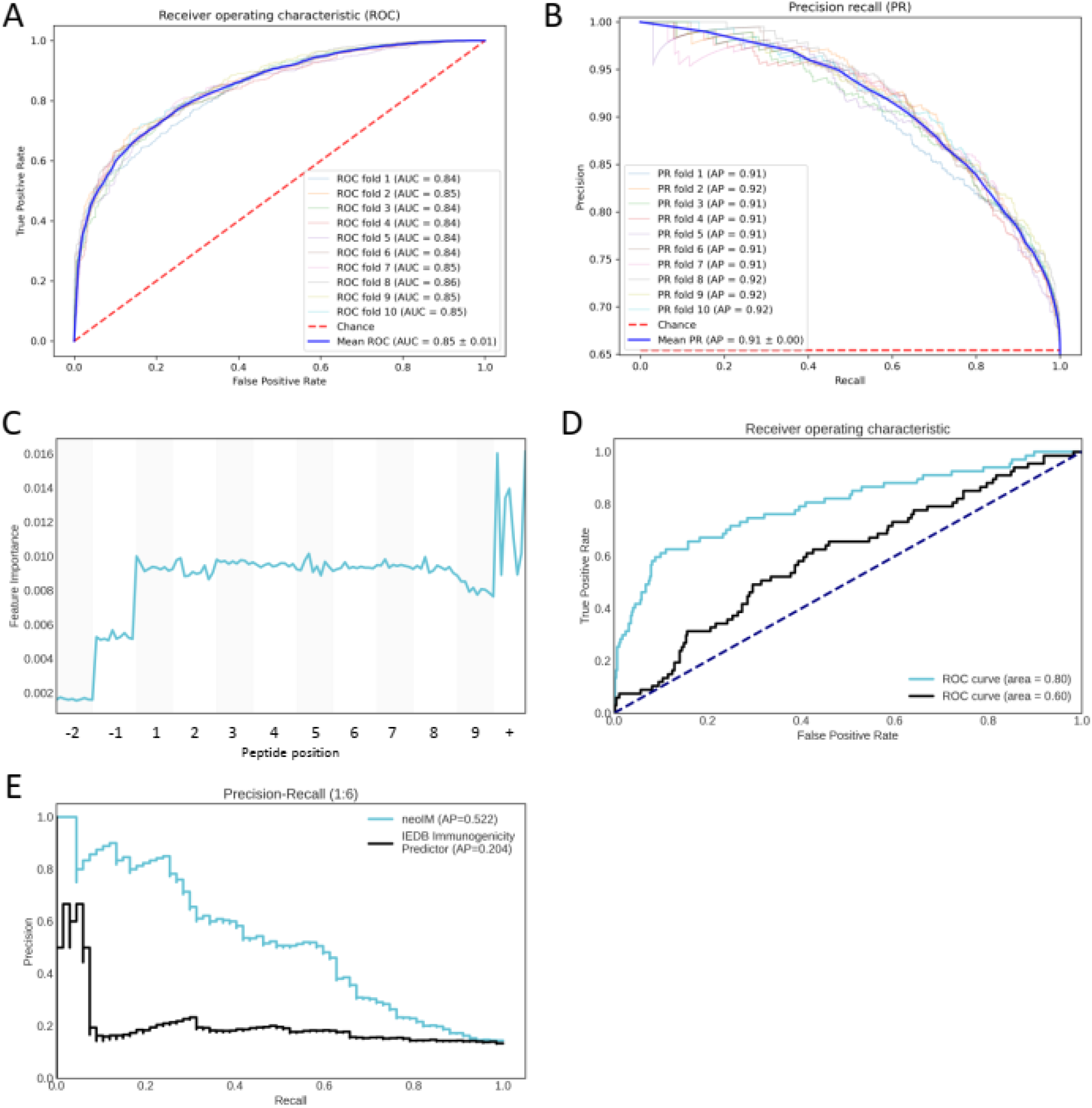
Panel A and B show the Receiver Operator Characteristic (ROC) (A) and Precision Recall (PR) (B) curves resulting from 10-fold cross-validation. Panel C shows the impact of features on model performance. Panel E and F show the ROC (E) and PR (F) curves resulting from benchmarking on an external dataset.

To determine which peptide positions, have the highest impact on model performance, we looked at feature importance (Figure 3C). Positions -2 and -1 are buffer positions that are imputed in peptides of shorter length and therefore, as expected, are of less importance. The last peptide position (position 9), which is a known anchor residue for MHC binding, also has a lower impact on model performance than the other peptide positions. As all peptides in the positive and negative training dataset are known MHC binding peptides, it is expected for an MHC anchor position to be less relevant. All other positions along the peptide sequence seem to be of similar importance to the model. The additional peptide-wide metrics (position +), display a higher importance than other positions, most likely due to them being an average of amino acid properties of the peptide. The top 10 most correlated amino acid properties per principal component (PC) are available in Supplementary Table 5.

Given that a length bias was identified in the training datasets, its effect on the performance of neoIM was investigated. To assess whether the algorithm is biased towards certain peptide lengths, the fractions of randomly selected human proteome peptides predicted to be immunogenic were compared across various peptide lengths. Within this framework, 9mers (13% positive) were less likely to be predicted as immunogenic than 10-(36%) or 11mers (41%) (p-value = 0.0, ANOVA). However, 10- and 11mers (32% and 28 %, respectively) were less likely to be presented than 9mers (39%) representing 32% and 28% of presented peptides versus 9mers representing 39% of presented peptides (Supplementary Figure 1C).

To investigate a potential bias of neoIM towards HLA-A*02:01-presented peptides, sets of randomly selected peptides from the human proteome, predicted to be presented across a wide range of HLA alleles (38 HLA-A, 52 HLA-B and 12 HLA-C alleles), were scored by neoIM. As shown in Supplementary Figure 1D, there is no significant difference between neoIM prediction ranges across HLA alleles (p-value = 0.27, ANOVA), with HLA-A*02-01 not presenting as an outlier and therefore showing no inherent bias of the algorithm towards HLA-A*02-01 presented peptides.

### Benchmark

To confirm its high performance, the neoIM model was further validated on an independent external dataset and compared to the only other comparable tool that is HLA agnostic and predicts immunogenicity based on the peptide sequence alone, the IEDB immunogenicity predictor^35^.

A validation dataset was constructed (Table 1) containing peptides that were all predicted to be presented and for which immunogenicity score was computed by both neoIM and the IEDB immunogenicity predictor. From the prediction scores, ROC curves and precision recall (PR) curves were constructed and are shown in Figure 3E and F. In the comparison, neoIM outperforms the IEDB immunogenicity predictor with an auROC of 0.79 vs 0.61 and an AP of 0.53 vs 0.23.

### In vitro validation results

ELISPOT validation experiments were performed using immune cells from three healthy donors on a set of 33 peptides derived from tumor-specific events, including SNVs (n=14), indels (n=7), gene fusions (n=5), retained introns (n=5) and long non-coding RNAs (n=2). All peptides were predicted to be presented on at least two donor HLAs to ensure peptide presentation and minimize the risk of false negative results.

Of the 33 peptides, seven peptides (21.2%) tested positive for at least one donor (Supplementary Table 6). Supplementary Figure 2A shows that peptides with a positive ELISPOT result (n = 7) had a significantly higher predicted immunogenicity score than peptides with a negative result (n = 25, p-value = 0.02). Using other methods, including MHC binding prediction alone and IEDB immunogenicity predictor, there is no significant difference in the scores for immunogenic and non-immunogenic peptides (Supplementary Figure 2B-D). This is also reflected in the ROC curve, where neoIM model is able to differentiate between immunogenic and non-immunogenic peptides with higher accuracy (Figure 4A). Furthermore, the neoIM score also shows the highest correlation, albeit not significant, with the strength of the detected immune response (Supplementary Figure 3, p-value = 0.17). The predicted strength of MHC binding (%Rank score) did not correlate with peptide immunogenicity. This shows that for peptides with a rank score below the binding threshold, the rank score should not be used to further stratify the peptides as immunogenic or not.

**Figure 4:**
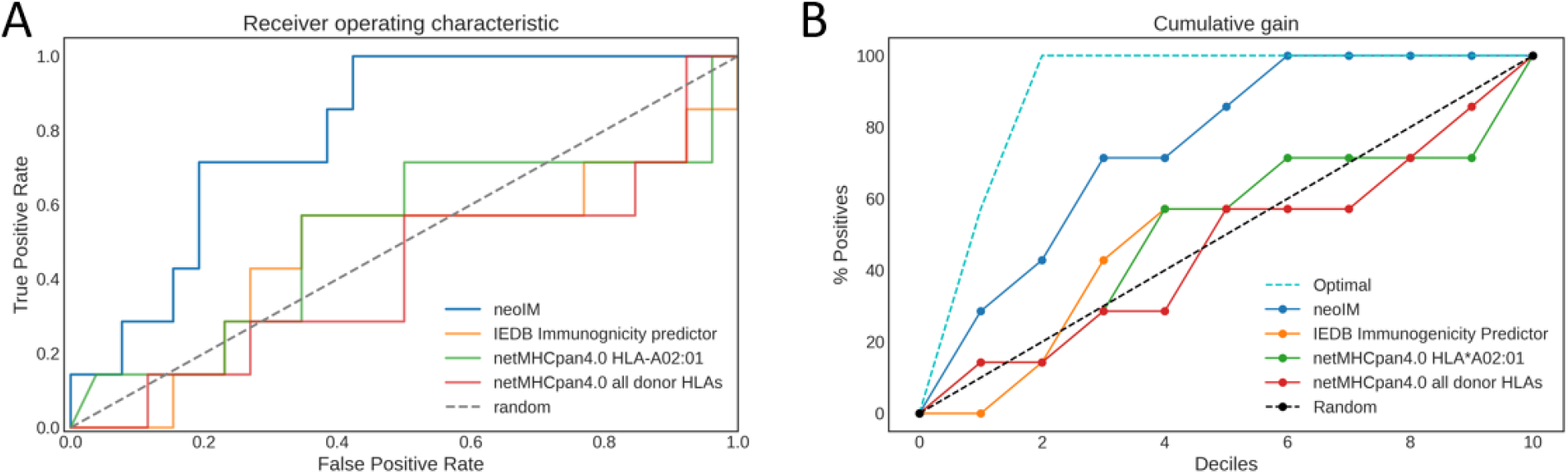
The Figure shows the correlation of different parameters used to predict an immunogenic reaction with ELISPOT results of 33 peptides which have all been predicted to be presented by netMHCpan4.0. Panel A shows a comparison of ROC curves of multiple different predictors, including the invention, netMHCpan4.0 and IEDB immunogenicity predictor. Panel B shows the lift curve of multiple different predictors, which plots the percentage of true positives (y-axis) in the top n deciles of ranked predictions.

The lift curve in Figure 4B shows the proportion of peptide immunogenicity scores ordered from highest to lowest (x-axis) in relation to the number of true positive predictions (y-axis). The curve shows that all positive peptides are represented in the top 55% of peptide immunogenicity scores, making it possible to discard 45% of all peptides without losing any true positives. This results in a significant increase of the true positive rate from 21% to 39% without losing sensitivity, substantially reducing the number of candidate neoantigens. Additionally, 71% of all immunogenic peptides are present in the top 30%, resulting in a further increase of the true positive rate to 50%.

## CPI Therapy

Next to PD-L1 expression, one of the most common clinical biomarkers for response to checkpoint inhibition therapy is tumor mutational burden (TMB)^43,44^. While high TMB is indeed generally correlated with long-term clinical benefit of immunotherapy, some tumors with low TMB still respond to and benefit from therapy. For these patients, TMB is insufficient as a clinical biomarker, and other biological parameters should be investigated to distinguish potential responders from non-responders.

In order to evaluate whether immunogenicity levels as predicted by neoIM could serve such a purpose, the number of immunogenic variants, the number of presented variants and the overall TMB^39^ were compared in a cohort of melanoma patients for which sequencing information as well as clinical outcome after CTLA-4 blockade was available.

Figure 5 shows Kaplan-Meier curves that estimate response rates based on the recorded duration of response to CPI therapy for various patient groups in the cohorts. In TMB low patients, overall TMB and the number of presented variants are not able to differentiate responders from non-responders (p-values = 0.94 and 0.77). An inverse trend can even be observed, wherein patients with a lower number of mutations display a higher response rate, however it is not conclusive. The number of immunogenic variants, in contrast, is able to significantly differentiate responding and non-responding patients (p-value = 0.04), with higher a higher number of immunogenic variants observed in responders. Therefore, the number of immunogenic variants can be used as a biomarker to identify patients with a low overall TMB that carry enough immunogenic mutations to benefit from immunotherapy.

**Figure 5:**
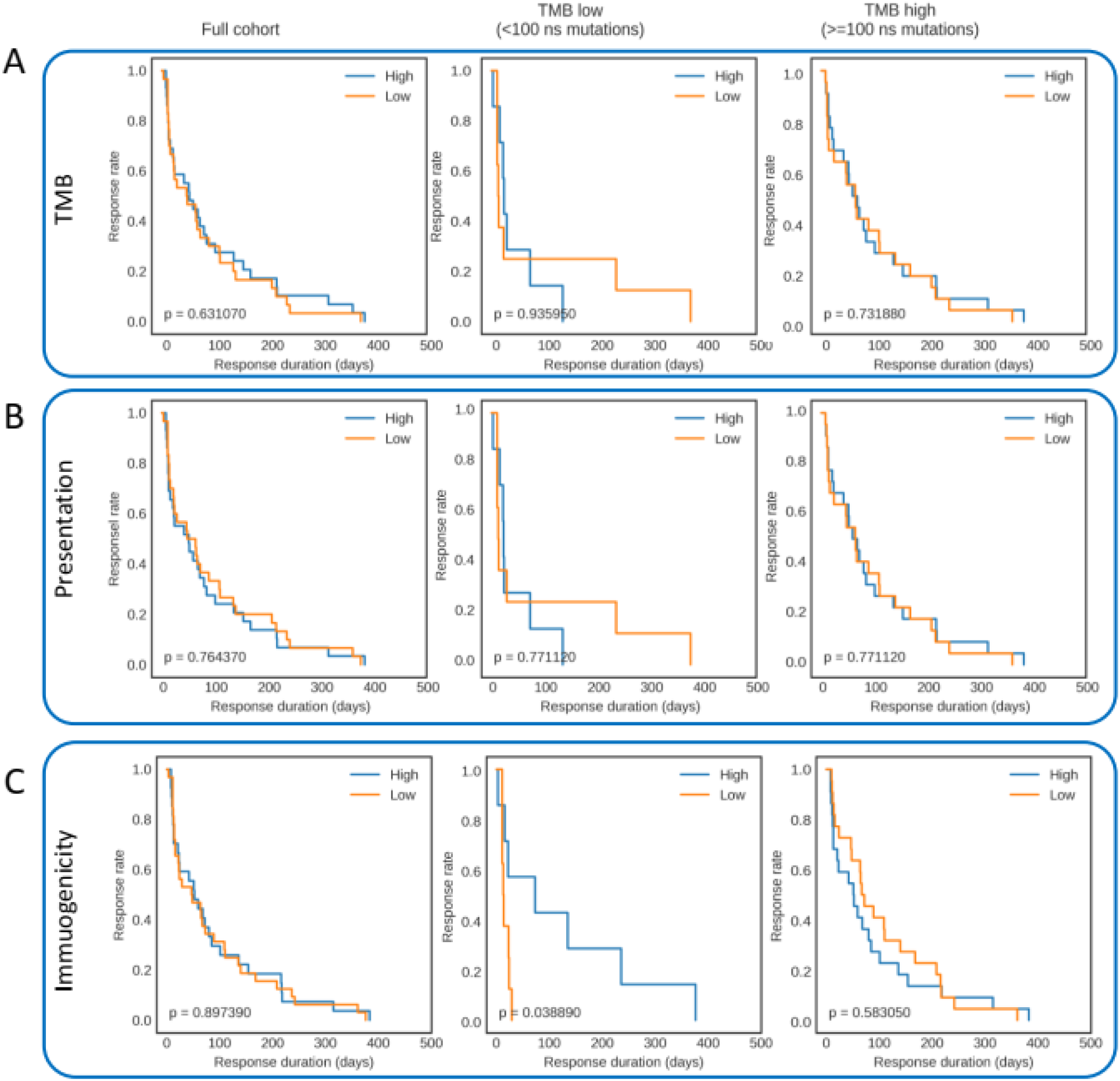
Kaplan-Meier plots showing the response rate (y-axis) versus the response duration (x-axis) of melanoma patients during CTLA-4 blockade. The survival curves show the patients divided into two groups, indicated as “High” (blue) or “Low” (orange), by the median number of mutations (A), number of presented mutations (B) or number of immunogenic mutations (C).

## Discussion

The quality of a machine learning model is highly dependent on its input, and negative data is just as important as positive data in building a training dataset that accurately represents the classes that should be predicted. As such, and with positive training data (i.e., peptides experimentally validated as being immunogenic) being largely available, efforts in establishing a correct training dataset for neoIM largely focused on curating an appropriate negative training dataset. The large differences in MHC polymorphisms and immune contexture between individuals, make it very challenging to obtain truly non-immunogenic peptides through experimental means. It is common to use randomly sampled (non-mutated) self-peptides as a negative background, which should not be recognized by T cells due to negative selection of T cells in the thymus. However, this may create a biased model that predicts whether a peptide is mutated or foreign rather than immunogenic.

A more appropriate background is a set of peptides originating from mutations, expressed and presented but non-immunogenic. In order to obtain a negative dataset that resembles this background, *in vivo* presented peptides identified by MS/MS were collected and peptides with mismatches to the human reference proteome were retained. These peptides are expected to originate from regions of genetic variation and thus should be more similar to the peptides of interest for immunogenicity prediction, peptides originating from mutations. We expect the majority of the peptides to stem from SNPs and only a very small fraction of them being potentially immunogenic as even for peptides that are actively prioritized only a fraction can be functionally validated as immunogenic^12^. Furthermore, germline mutation rates are much lower than somatic mutation rates and both display distinct mutation spectra, highlighting the importance of protecting the integrity of the germline genome^45^. Thus, it could be hypothesized that germline mutations are in general less disruptive to the genome which could mean that they are also less likely to be immunogenic.

When comparing the peptides in the positive dataset, which have been actively selected for immunogenicity testing, with the peptides in negative dataset, which consist of MS detected ligands, an unequal length distribution can be observed. More precisely, there is a higher fraction of longer peptides in the positive dataset, which could indicate a potential bias trained into the model. The model indeed outputs on average a higher immunogenicity score for longer peptides. Further research is needed to determine whether this is due to a biased model or if there is an actual relationship between peptide length and immunogenicity^12,46^.

The observed drop of the average precision of neoIM when validated on the external dataset as compared to cross validation, is most likely due to false negative peptides in the external dataset. The classification of peptides as immunogenic or non-immunogenic in this dataset is based on *in vitro* validation experiments. However, several peptides classified as non-immunogenic in the external datasets were found to be immunogenic in other experiments, as recorded in the IEDB database. While the known false negative peptides were filtered out of the dataset before performing the validation experiment, it is likely that a number of false negatives still remained. These peptides could be responsible for the drop in precision in our model and highlight the difficulty in defining truly non-immunogenic peptides.

By analyzing differential amino acid usage of peptides in the positive dataset against the background of peptides in the negative dataset, some features of immunogenic peptides could be characterized. In general, immunogenic peptides were found to be enriched in hydrophobic amino acids, neutrally charged amino acids, large amino acids side chains and amino acids containing sulfur, aromatic or aliphatic groups. Hydrophobicity at MHC and TCR contact residues has been described before to be a predictor for peptide immunogenicity, as well as large and aromatic residues ^22,23,35^. However, the role of electrical charge and the role of sulfur containing residues, which may contribute to overall complex stability, is less documented^47^.

*In vitro* validation experiments demonstrate that neoIM is capable of distinguishing between immunogenic and non-immunogenic peptides with high accuracy. Importantly, even though all peptides were predicted to be presented, and were introduced at a high concentration in the assay, thus maximizing their chance of MHC binding, only a minority of them was able to elicit an immune response in healthy donors’ cells, highlighting the intrinsic difference in immunogenicity between epitopes. However, the data also shows that peptides can be immunogenic and stimulate T-cells in donors without a pre-existing response, which highlights the importance of immunotherapies that re-engage a patient’s immune system. Moreover, a clear correlation was found between the neoIM immunogenicity score and the T-cell response rate allowing to define an optimal immunogenicity threshold thereby lowering the false positive rate.

By using neoIM to rate the immunogenic potential of mutations, a subset of melanoma patients with low overall TMB but a high fraction of immunogenic variants has been identified that is more likely to respond to immunotherapy than patients with a low TMB and low fraction of immunogenic peptides. This is in line with other recent studies that have shown a stronger correlation between response to immune checkpoint inhibition with the number of immunogenic variants rather than mutational burden^12,23,48^. For patients with high TMB, in comparison, further classification into low- and high-immunogenicity patients seemed to have little benefit, most likely due to a higher base-level of immunogenicity across the entire population derived from a statistically higher number of immunogenic epitopes by random occurrence. In light of these results, neoIM can be a powerful tool both in improving target selection for personalized immunotherapies, especially in patients with low TMB, as well as in offering immunogenic peptide load as an improved biomarker for patient survival and response to therapy. However, further studies including more patients with low TMB and other cancer histotypes are necessary to determine if variant immunogenicity can be a universal biomarker in low TMB tumors.

In this study, peptide immunogenicity prediction was solely based on physicochemical features. However, this only represents a part of what drives an immune response, and as such, other important features, e.g., gene expression or variant allele frequency, could also be taken into consideration in future versions of the algorithm presented here. Furthermore, we are continually working on improving and expanding the training dataset. With more data becoming available in the future for instance, some potential issues of unbalance in the dataset, i.e., differences in size and HLA distribution in positive and negative dataset can be addressed. Importantly, the methodology described here is also applicable to peptides presented on MHC class II, although there is less experimental data available, and work is underway to develop a version of neoIM for longer peptides presented on MHC class II. This will lead to a more complete picture of a patient’s immune response and can hopefully further improve therapy outcomes.

In conclusion, we demonstrate that neoIM is an attractive option for immunogenicity prediction and neoantigen screening. By using a training dataset only containing peptides that are known to be presented, the model focuses on T cell recognition rather than MHC presentation. This makes neoIM an HLA-agnostic model, and therefore an easily applicable tool to predict immunogenicity for any presented peptide. The only other comparable tool, the IEDB Immunogenicity Predictor, is significantly outperformed by neoIM. Because neoIM was developed for the purpose of scoring the immunogenic potential of cell-surface presented peptides, peptides that are scored by neoIM should therefore be peptides that are predicted or known to be presented on MHC molecules. In the context of neoantigen prediction pipelines, neoIM would optimally be used as an additional module to variant detection, expression analysis and presentation prediction. Prioritizing neoepitopes with high predicted immunogenicity by neoIM increases the likelihood of selecting truly immunogenic peptides. This reduces the number of peptides necessary to screen when selecting target epitopes for immunotherapy and lead to a time and cost reduction. Furthermore, it can be a valuable tool in prioritizing actionable neoantigens in the context of personalized therapy where it might not be feasible to experimentally validate all candidate epitopes.

## Supplementary

**Supplementary Figure 1:**
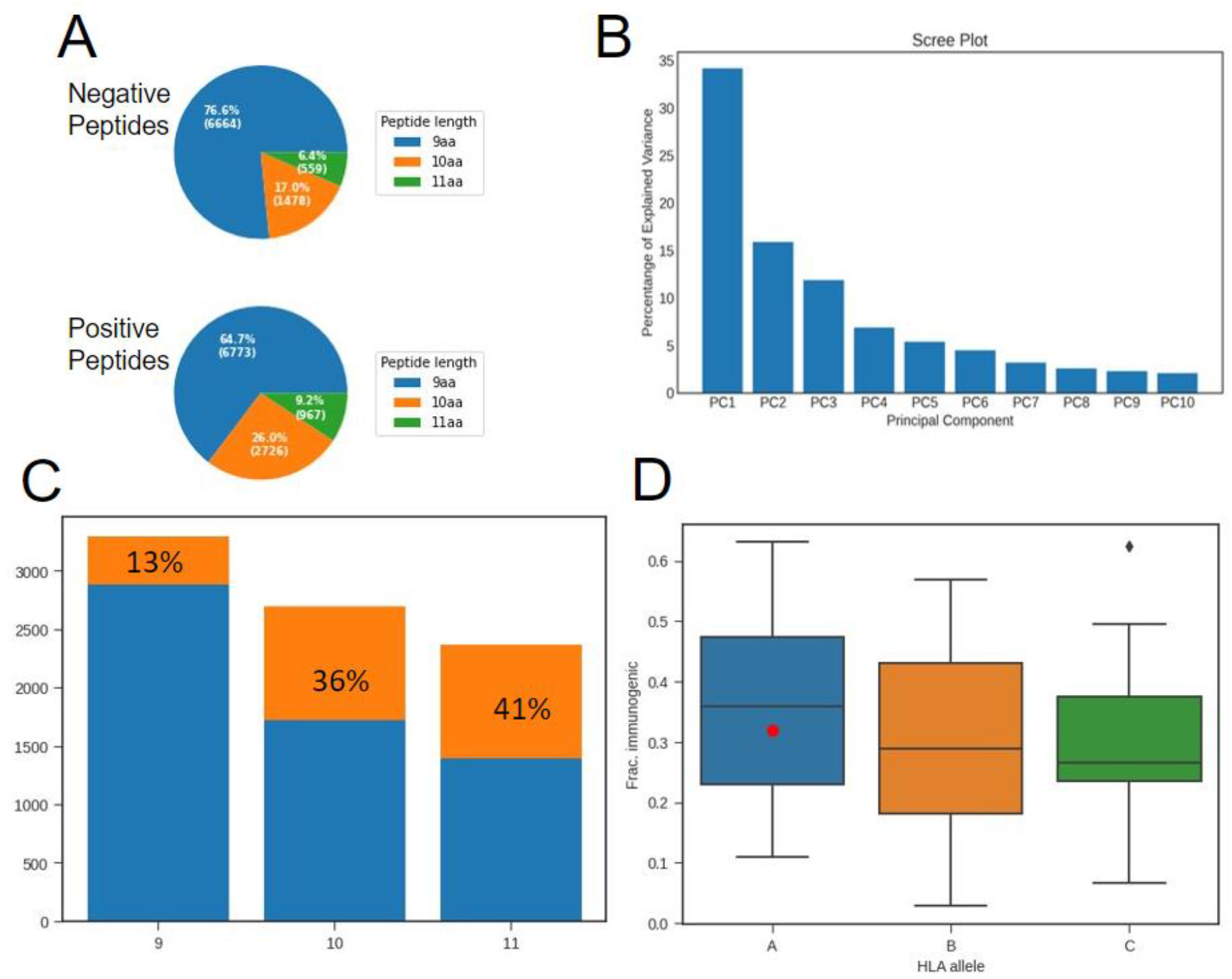
Panel A shows the distribution of peptide lengths for the positive and negative training dataset. Panel B shows the percentage of explained variance by each principal component for amino acid properties used for peptide encoding. Panel C shows the number of randomly selected peptides that were considered binders (out of 25000 peptides) with the fraction of peptides determined to be immunogenic by neoIM colored in orange. Panel D shows the fraction of peptides determined to be immunogenic when presented by various HLA alleles, grouped by HLA-A, HLA-B and HLA-C, with the value for HLA-A*02:01 shown as a red dot.

**Supplementary Figure 2:**
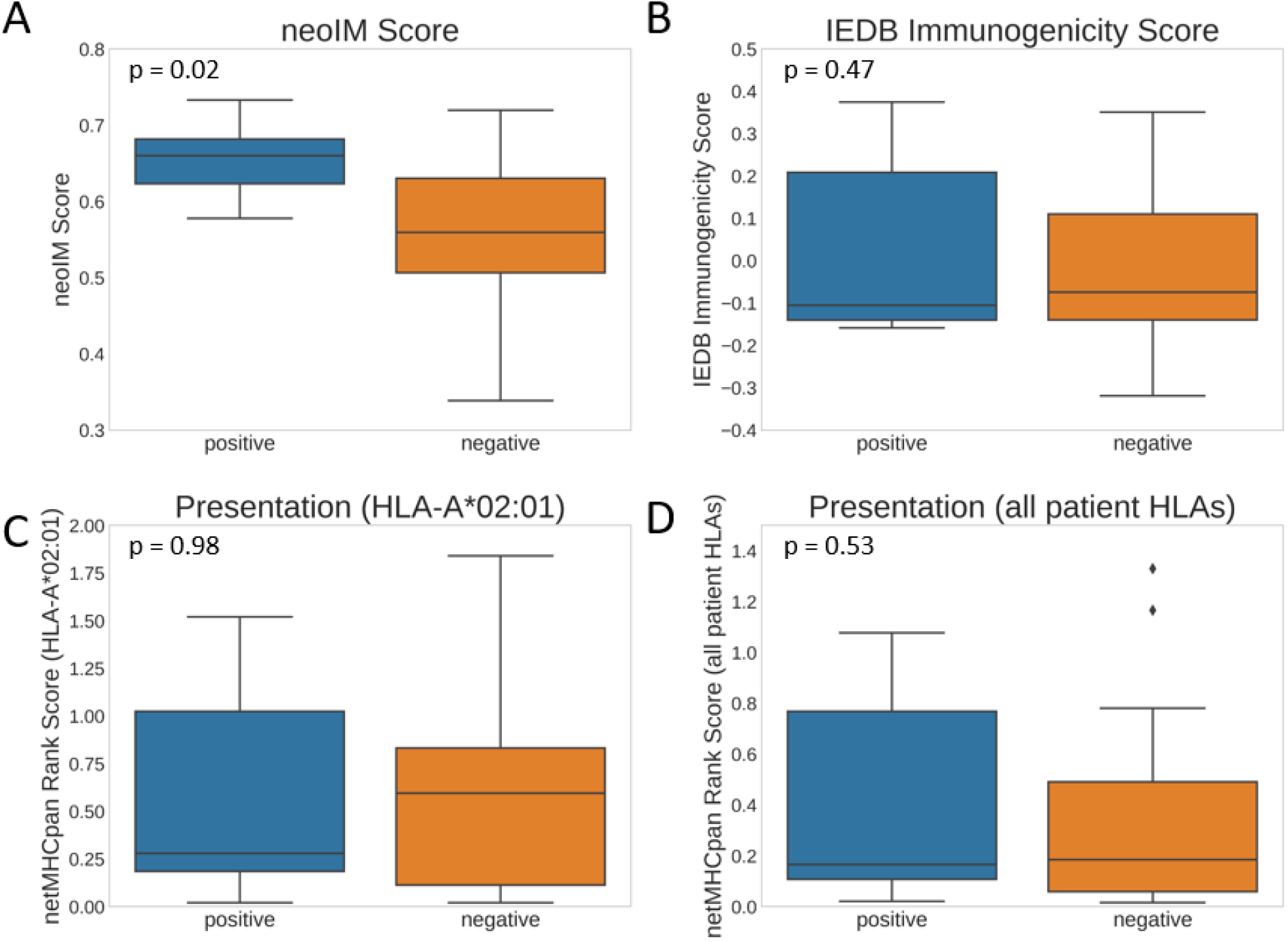
This figure shows the scores for neoIM (A), IEDB Immunogenicity predictor (B), netMHCpan rank score (HLA-A*02:01, C) and netMHCpan rank score (lowest of all patient HLA alleles, D) for all peptides evaluated as positive or negative in the *in vitro* validation dataset.

**Supplementary Figure 3:**
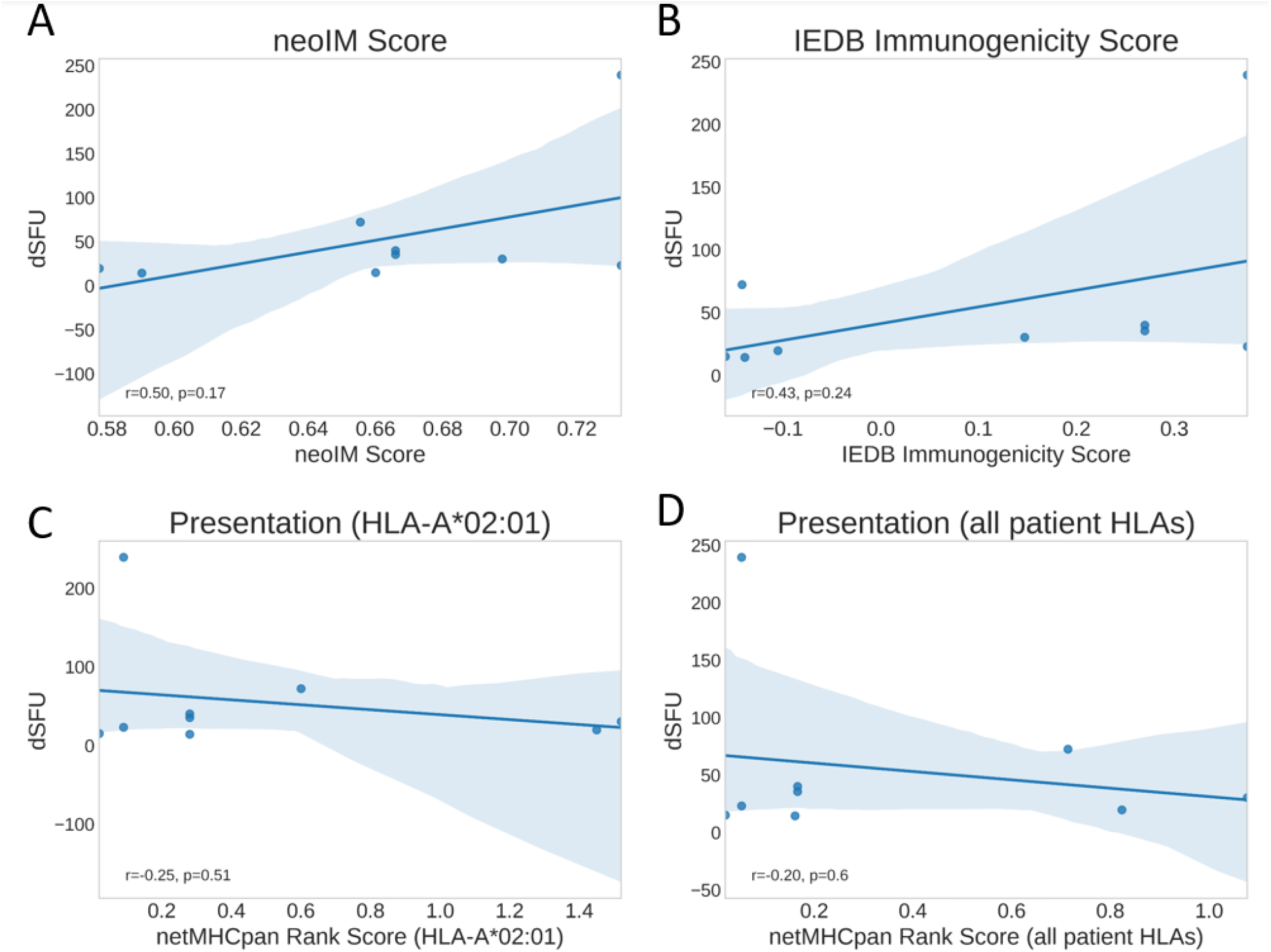
This figure shows the Pearson correlation for neoIM (A), IEDB Immunogenicity predictor (B), netMHCpan rank score (HLA-A*02:01, C) and netMHCpan rank score (lowest of all patient HLA alleles, D) with the strength of the immune response in Spot Forming Units (SFU) between the test condition and reference condition (dSFU) for all peptides evaluated as positive in the *in vitro* validation dataset.

